# Multiscale cardiac imaging to capture the whole heart and its internal cellular architecture, with applications to congenital heart disease

**DOI:** 10.1101/2020.04.22.055418

**Authors:** Graham Rykiel, Claudia S. López, Jessica L. Riesterer, Ian Fries, Sanika Deosthali, Katherine Courchaine, Alina Maloyan, Kent Thornburg, Sandra Rugonyi

## Abstract

Efficient cardiac pumping depends on the morphological structure of the heart, but also on its sub-cellular (ultrastructural) architecture, which enables cardiac contraction. In cases of congenital heart defects, localized sub-cellular disruptions in architecture that increase the risk of heart failure are only starting to be discovered. This is in part due to a lack of technologies that can image the three dimensional (3D) heart structure, assessing malformations; and its ultrastructure, assessing disruptions. We present here a multiscale, correlative imaging procedure that achieves high-resolution images of the whole heart, using 3D micro-computed tomography (micro-CT); and its ultrastructure, using 3D scanning electron microscopy (SEM). This combination of technologies has not been possible before in imaging the same cardiac sample due to the heart large size, even when studying small fetal and neonatal animal models (~5×5×5mm^3^). Here, we achieved uniform fixation and staining of the whole heart, without losing ultrastructural preservation (at the nm resolution range). Our approach enables multiscale studies of cardiac architecture in models of congenital heart disease and beyond.

## Introduction

Congenital heart disease (CHD), which manifests as a morphologically defective heart, affects about 1% of newborn babies, and remains the primary cause of non-infectious children mortality in the developed world^1,2^. While CHD mortality rates have been dramatically reduced in recent years thanks to advances in surgical practice and interventional technologies^1,3^, CHD patients continue to be at an increased risk of developing heart failure at a much younger age than the general population^4^. Despite early indicators of success, heart failure continues to take the lives of young children with CHD: 10 to 25% of newborns with a critical heart defect do not survive the first year, and 44% do not survive to 18 years of age^5,6^ This unfortunate trend points to cardiac deficiencies in CHD that are not yet understood^7^.

While the structural (morphological or “geometrical”) characteristics of heart malformations have been extensively studied, it is largely unknown whether cardiac cells from malformed hearts are normal or to what extent they are compromised. Recent studies reveal an abnormal orientation of myocardial cells (the heart muscle cells) within CHD hearts^8–10^. Myocardial cells are elongated, cylindrical-like cells, that contract along their long axis. In a normal heart, myocardial cells arrange in sheet-like layers with their long axes in parallel to each other forming an elliptical pattern^11,12^. Newly developed contrast episcopic microscopy and synchro micro-CT imaging techniques, enable non-destructive analyses of banked human fetal and neonatal hearts with CHD.^9,10^ These studies are revealing myocardial disarray in CHD (with respect to their normal counterparts) that very likely affect cardiac function after surgical repair, and that have been ignored when planning treatment strategies for CHD patients. In addition to changes in the myocardial organization, the sub-cellular contraction machinery of myocardial cells (e.g the myofibrils that contract the cell; and the mitochondria that provide energy for contraction) may also be compromised in CHD, affecting heart function. The extent to which the cells of malformed hearts exhibit deficiencies is unknown^8,10,13^. This is in part due to limitations of existing technologies that have not achieved precise multiscale mapping to decipher the association between structural and cellular deficiencies in the heart and beyond.

We describe here a novel, correlative multiscale imaging procedure that combines imaging of whole heart morphology and its sub-cellular organization (ultrastructural architecture). Our multiscale procedure uses micro computed tomography (micro-CT) imaging to capture heart morphology at micrometer resolution, and scanning electron microscopy (SEM) to capture cardiac tissue ultrastructure at nanometer resolution. Current SEM technologies allow for threedimensional (3D) imaging of sub-cellular architecture, enabling reconstruction and quantification of ultrastructural features within a tissue volume^14–16^. Among 3D SEM methods, we have selected serial block-face SEM (SBF-SEM) for ultrastructural imaging, as it allows 3D imaging of relatively large volumes (sample size 40×60×40 μm^3^). The methodology we present herein improves upon previous protocols by achieving uniform staining of a relatively large heart sample (3-4 mm wide, 5-6 mm long), circumventing micro-CT x-ray penetration issues, and allowing sample screening and selection prior to full sample preparation. Our multiscale imaging, further, enables mapping of structural and ultrastructural heart features.

As proof of concept, we applied our developed multiscale imaging procedure to two embryonic chick hearts. These hearts were collected at stages corresponding to about 5-6 months of human fetal development, when the heart is already formed but maturing in preparation for birth/hatching. We imaged: 1) a control heart with no structural defects; and 2) a heart with tetralogy of Fallot (TOF), a combination of structural heart malformations found in humans^17,18^. Our results suggest differences in the ultrastructure of these two hearts, emphasizing the need for a multiscale approach to deepen our understanding of CHD and enable the development of effective strategies to combat heart failure in CHD.

## Results

### Overview of multiscale imaging procedure

To achieve multiscale imaging we followed a four-step protocol (see **Figure 1**; details in Methods). Briefly, in **Step 1** the heart was excised, homogeneously fixed and stained for microCT. Initial staining followed a modified ferrocyanide-reduced osmium–thiocarbohydrazide-osmium (ROTO) protocol^19–21^ typically used for electron microscopy (EM) sample preparation. Three-dimensional micro-CT images (10 μm resolution) confirmed uniform ROTO staining of the whole heart and provided morphological cardiac details. At this time the heart samples were stored until further processing, enabling selection of specific samples for full processing based on micro-CT scans. **Step 2** finalized the preparation of the whole heart for SBF-SEM by poststaining the hearts with uranyl acetate and lead aspartate and then embedding them in a resin block. Uniform staining was confirmed on a semithin (250 nm) section of the block, which also determined ultrastructural quality and enabled registration to micro-CT images. In **Step 3**, a slab of the sample was cut and sectioned around a specific region of interest (ROI, ~500×500 μm^2^) from which sub-ROIs for 3D SBF-SEM imaging (~ 40×60 μm^2^) were selected. In **Step 4**, 3D SBF-SEM datasets were acquired (10 nm lateral resolution and 40 nm depth resolution).

**Figure 1:**
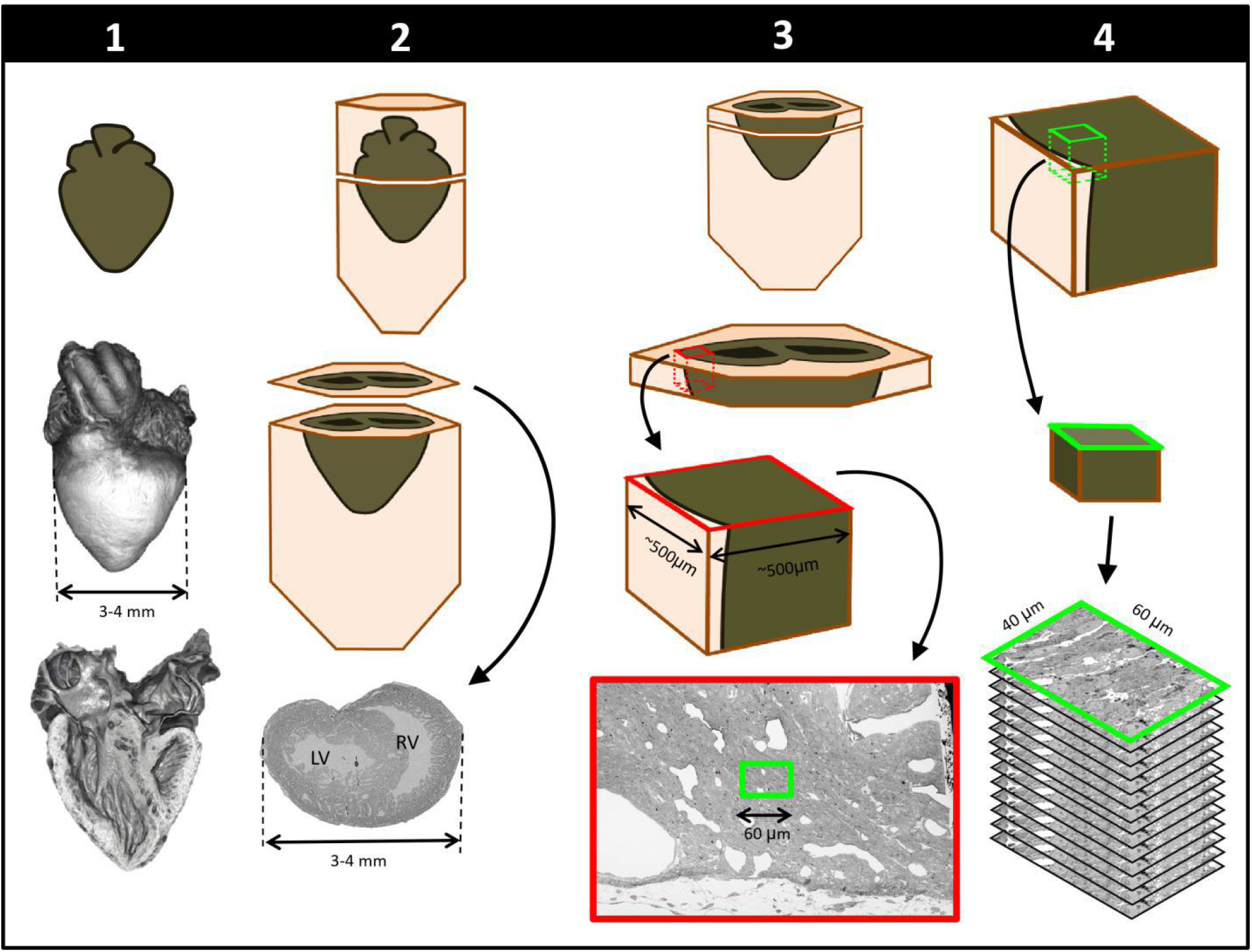
Schematics of steps performed to achieve cardiac multiscale imaging, which yields both 3D whole-heart images and 3D ultrastructural images from the same heart. Columns depict the four steps employed. In **Step 1**, the heart is post-fixed with osmium tetroxide to provide contrast for 3D micro-CT images of the whole heart (middle row). Digital sections of the microCT scans (bottom row) reveal the heart’s interior and allow for cardiac phenotyping and assessment of stain penetration. In **Step 2**, contrast staining is finalized and the resin block in which the heart is embedded is sectioned to reach a plane of interest (bottom row). In **Step 3**, after cutting a slab of the sample, a region of interest (ROI) is sectioned from the slab, mounted, and then scanned by SEM backscattered imaging methods to aid in the selection of sub-ROIs (for example, the sub-ROI highlighted in green). In **Step 4**, the selected sub-ROI 3D SBF-SEM images are acquired by progressively sectioning and imaging 40nm thick layers.

### Whole heart imaging

We obtained 3D micro-CT images of the whole heart, featuring both external and internal structures at 10 μm resolution (**Figure 2**; **Step 1** in **Figure 1**). Despite the relatively large dimensions of the heart (5-6 mm long; 3-4 mm wide), tissue contrast was uniform across the heart walls and septae and allowed us to visualize the microstructural details of the heart chambers, valves, and great arteries.

**Figure 2:**
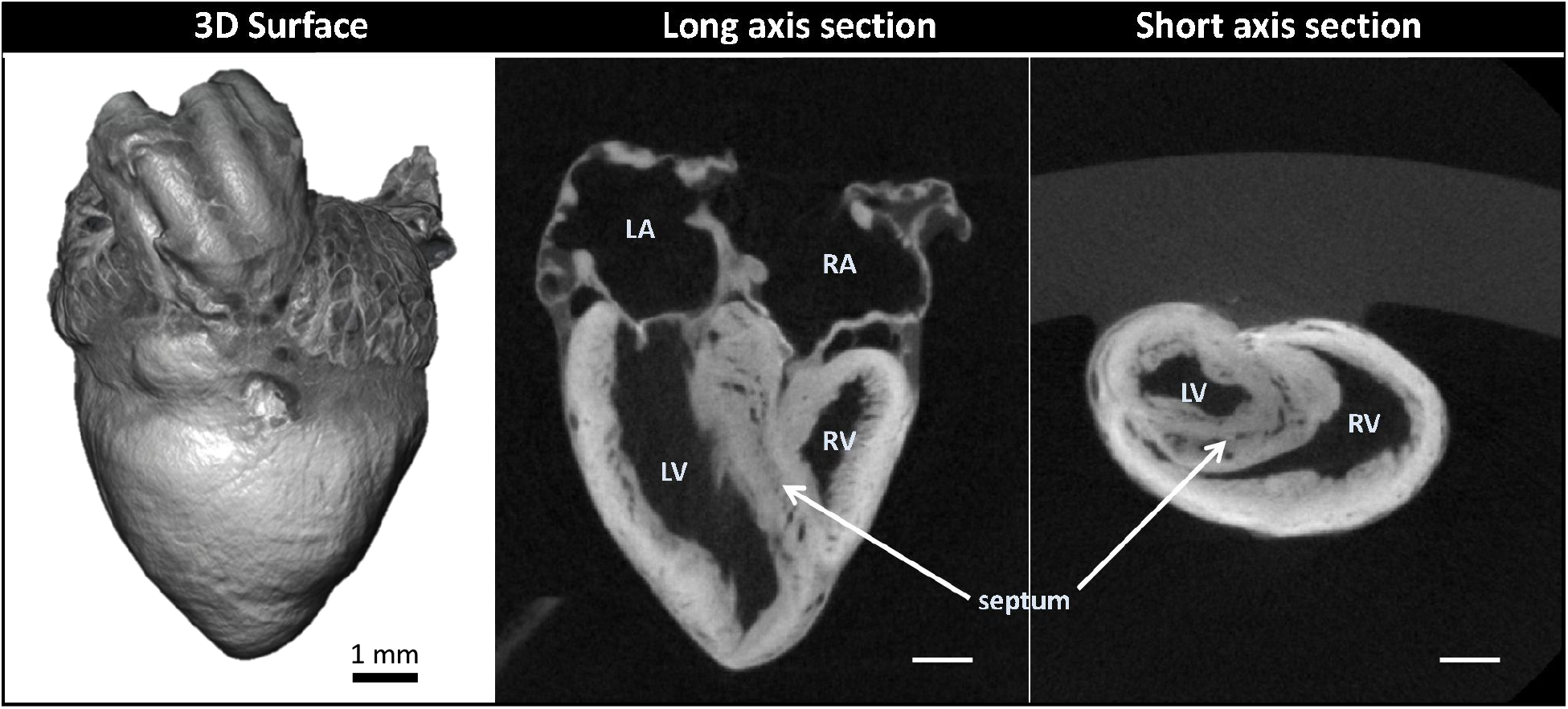
Micro-computed tomography (micro-CT) images of a chicken embryo normal heart. The contrast was achieved by following **Step 1** of our correlative multiscale imaging protocol (**Figure 1**). **From left to right:** External 3D surface of the heart; cardiac section along the heart’s long axis (coronal section); cardiac section across the heart’s short axis (transverse section). Cardiac sections show uniform staining across cardiac walls, and reveal the heart internal and external microstructure. LA: left atrium; LV: left ventricle; RA: right atrium; RV: right ventricle. Scale bars 1mm.

### Cardiac structure analysis from 3D micro-CT images

Micro-CT images were used to explore the structural characteristics of several chick hearts, to select two hearts for subsequent SBF-SEM imaging and analysis. We selected: 1) a normal heart; and 2) a heart exhibiting TOF malformation. In a normal heart, blood in the left ventricle (LV) and right ventricle (RV) is separated by an interventricular septum; the pulmonary valve and pulmonary artery connect to the RV, which pumps blood to the lungs; and the aortic valve and aorta connect to the LV, which pumps blood to the body. TOF is characterized by a combination of 4 defects: i) ventricular septal defect, which is a hole in the interventricular septum; ii) overriding aorta, a change in the position of the aorta such that it sits in the middle of the two ventricles, on top of the ventricular septal defect; iii) pulmonary stenosis or atresia, a narrowing or closure of the pulmonary artery or pulmonary valve; and iv) RV hypertrophy, a thickening and enlargement of the RV wall. RV hypertrophy in TOF, however, develops over time as the stenosis of the pulmonary artery increases pressure in the RV after birth^17^ and was not present in the heart examined in this study (see **Figure 3** for a comparison of the selected normal and TOF hearts). The TOF heart analyzed here featured supravalvular pulmonary stenosis, a ventricular septal defect, and an overriding aorta. The right ventricle was enlarged and thin-walled compared to the control heart (**Figure 3**). Further, the TOF heart was missing the right branch of the pulmonary artery. In humans, this rare condition, called unilateral absence of a pulmonary artery, is known to occur in conjunction with TOF or cardiac septal defects^22^.

**Figure 3:**
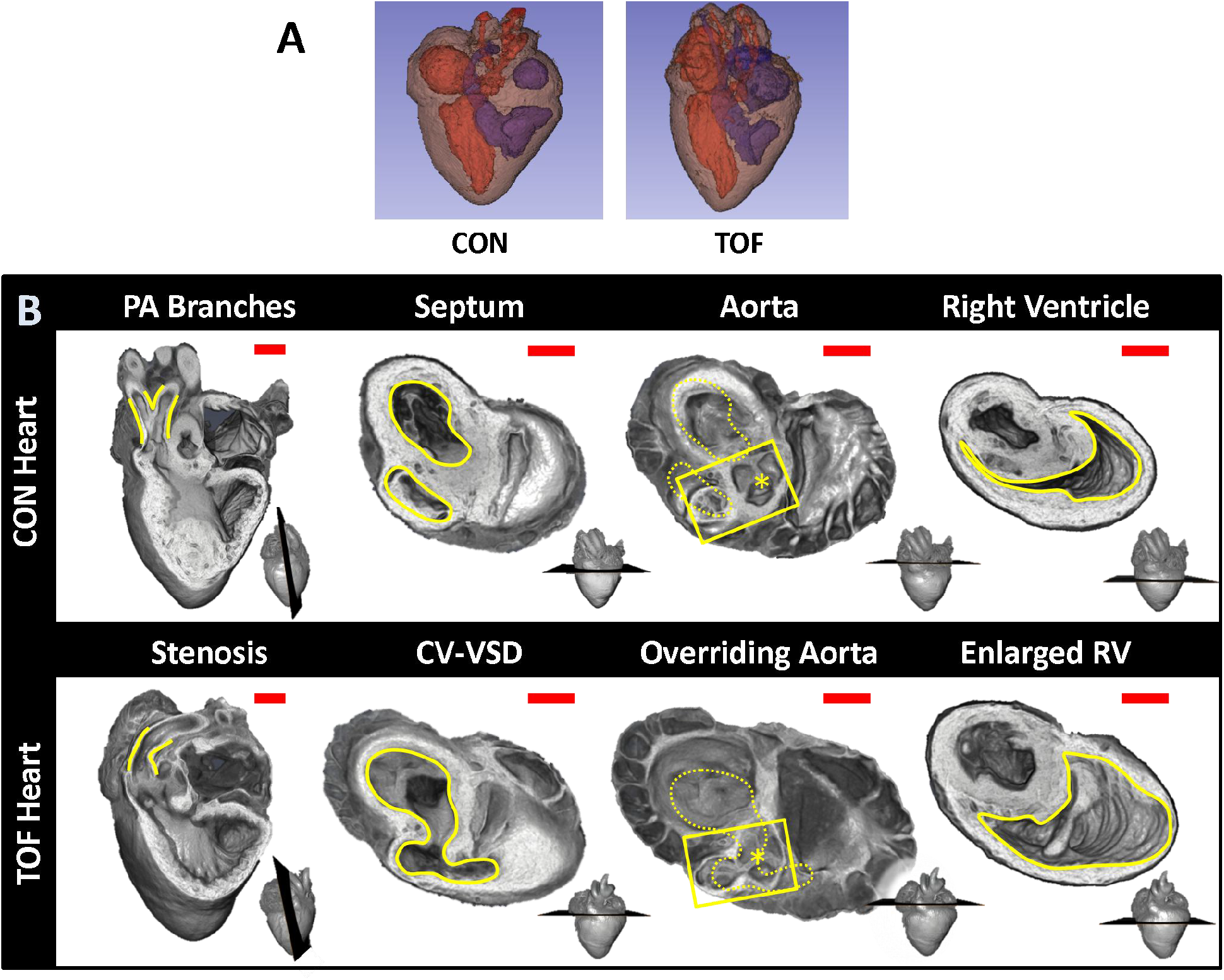
Comparison of micro-CT images of the two hearts selected for this study. **(A)** Segmentations showing the heart morphology for the normal, control (CON) heart and the TOF heart. Red: lumen of the left atrium and ventricle as well as aorta; Blue: lumen of the right atrium and ventricle as well as pulmonary artery; Brown: heart tissue. **(B)** Detailed comparison of the two hearts. Each column compares a cardiac feature (highlighted in yellow) between the hearts. The position of the plane along which the tissue was cut for display is shown at the bottom right of each image. From left to right: **Pulmonary artery (PA) branches:** On the control heart, the PA is bifurcated (yellow lines) whereas the left branch of the PA is absent in the TOF heart. The remaining PA of the TOF heart exhibits supravalvular stenosis (yellow lines). **Septum:** The ventricles (yellow lines) in the control heart are discrete, separated by an intact interventricular septum, whereas the TOF heart shows a conoventricular septal defect (CV-VSD). **Aorta:** In the control heart the aortic valve (asterisk) is connected to the left ventricle, whereas in the TOF heart the aortic valve is positioned directly over the VSD. Dotted yellow lines show the position of yellow lines in the septum column. **Right Ventricle (RV):** The RV (yellow lines) is significantly larger in the TOF heart than in the control heart, and features thinner walls. Scale bars = 1mm.

### Cardiac ultrastructural imaging

We chose to characterize the ultrastructure architecture of the selected hearts at approximately the transverse section at which the heart width (from LV to RV wall) is maximal, also referred to as the equatorial plane. Images of semithin cross-sections for each heart (**Figure 4**, top row; corresponding to **Step 2** in **Figure 1**) show that staining was uniform, indicating successful stain penetration through the heart tissues. Please note that the transverse section of the TOF heart was below the ventricular septal defect and thus exhibits a continuous septum.

**Figure 4:**
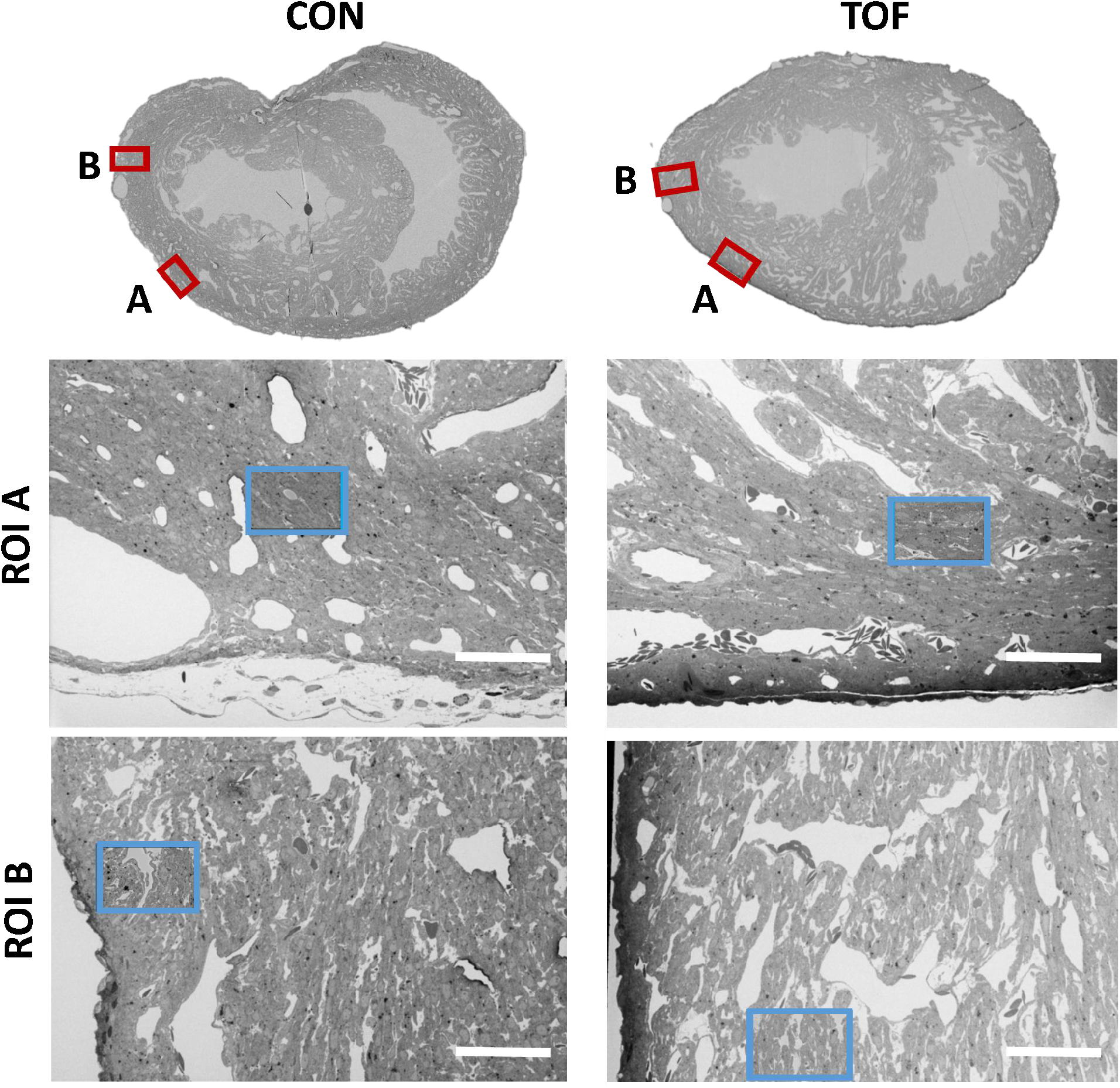
SBF-SEM images of the control (CON, left) and TOF (right) hearts, indicating the location of imaging. Overview scans from semithin transverse sections of each heart (**top row**) were used to confirm uniform stain penetration and the location of 2 ROIs per heart (ROI A and ROI B) further analyzed (red boxes). Semithin sections are not to scale. Blood vessels and trabeculae served as landmarks for accurate positioning of the ROIs and sub-ROIs. The sub-ROIs corresponding to ROI A (**second row**), were fully segmented (nuclei, extracellular space, myofibrils, and mitochondria). However, the sub-ROIs corresponding to ROI B (**third row**) were only partially segmented (only a fraction of the images in the stack were segmented) for quantification purposes. Scale bars 60 microns.

Using the semithin section SEM images, ROIs from the hearts were selected. For this study, we selected two ROIs within the heart LV, denoted by ROI A and ROI B. After cutting the sample, the ROIs were first imaged with SBF-SEM at low resolution (65-80 nm lateral resolution; **Step 3** in **Figure 1**), from which sub-ROIs were further selected (see **Figure 4**). These sub-ROIs (~40×60 μm^2^) were imaged at 10 nm lateral resolution, and we acquired 800-1000 images in depth (with ~40nm of depth distance, thus 32 to 40 μm in depth). These high-resolution images showed the conservation of ultrastructural features (see **Figure 5**). Images exhibited continuous nuclear membranes, intact mitochondria, and defined myofibrils, indicating that we achieved both appropriate and uniform fixation and staining of the hearts with our protocol.

**Figure 5:**
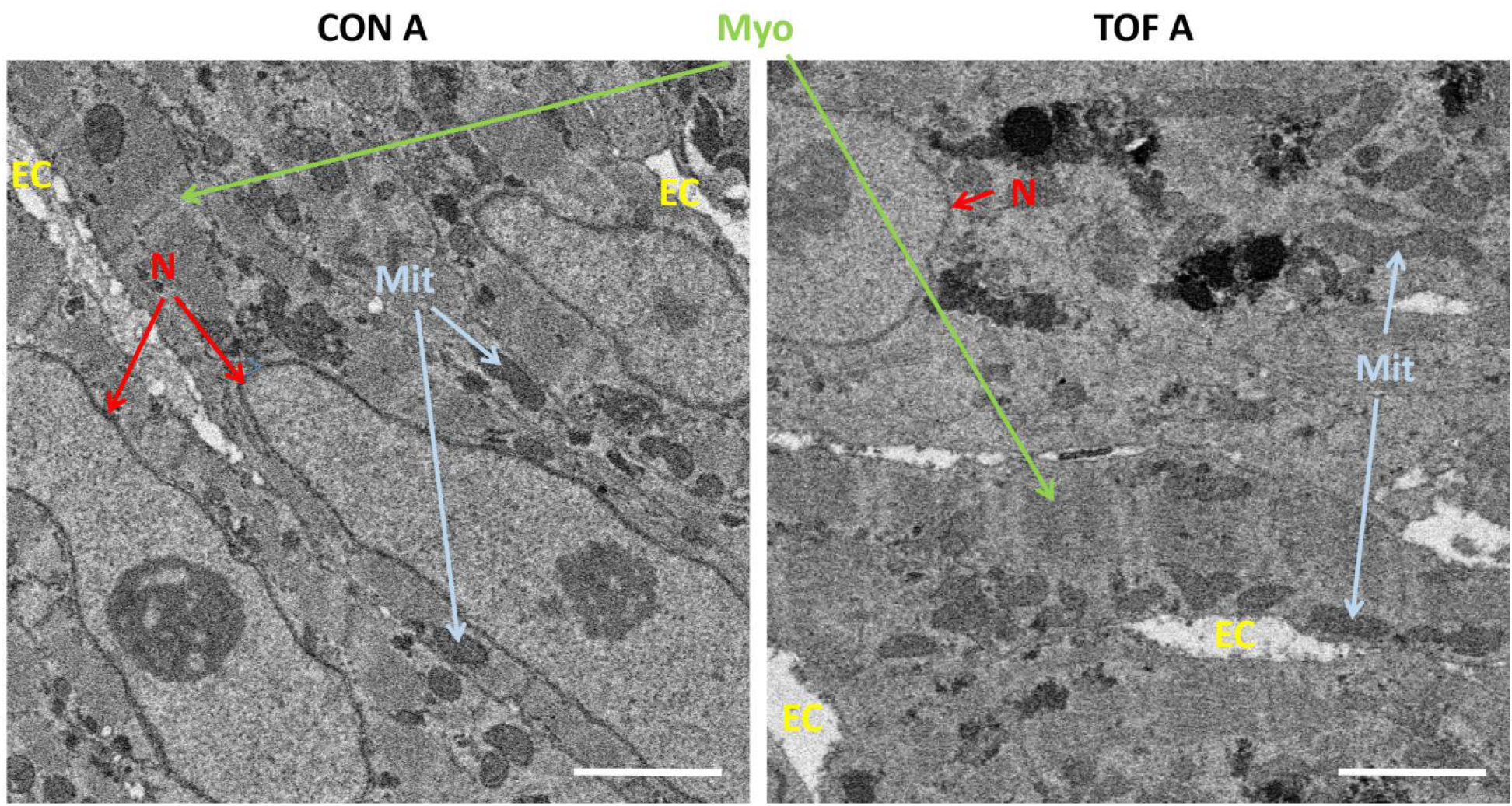
Detail of high-resolution SBF-SEM images obtained. The pictures depict small regions within the selected sub-ROIs from region A of the control (CON) and TOF hearts. Nuclear membranes (N) are depicted, as well as myofibrils (Myo) and mitochondria (Mit). Finally, the extracellular space (EC) is also visible. Scale bars 2 microns.

### 3D SBF-SEM image reconstruction

SBF-SEM image stacks provided 3D volumetric recontructions of sub-ROIs. While volumetric image resolution was not isotropic (10 nm lateral resolution versus 40 nm depth resolution), ultrastructural features could be visualized from any angle of view within the reconstructed images (see **Figure 6**). Thus SBF-SEM images allowed us to visualize the orientation and organization of nuclei, myofibrils, and mitochondria (among other features) in heart tissue samples.

**Figure 6:**
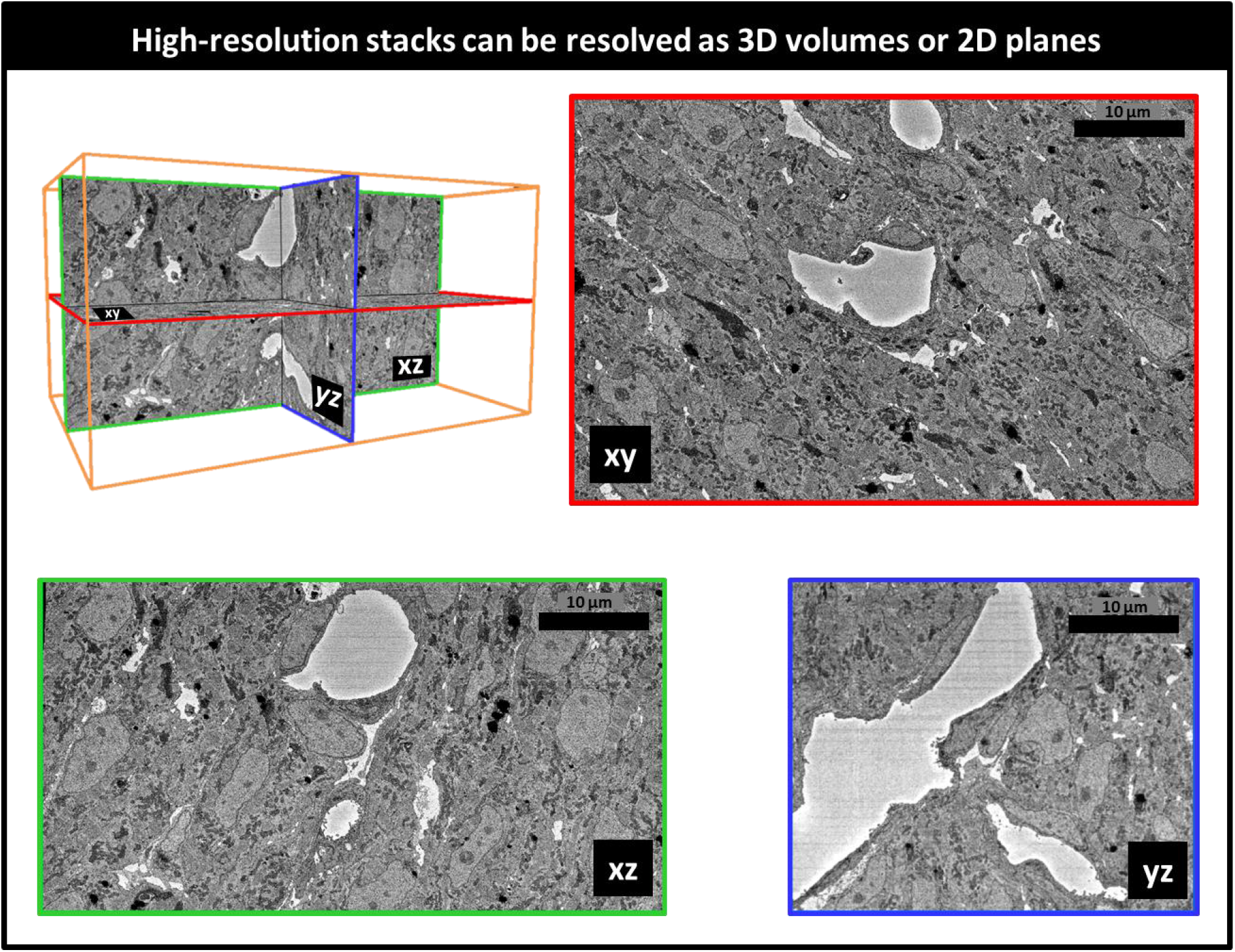
Example 3D reconstruction of SBF-SEM image stack acquired from the control heart. **xy** is the imaging plane, acquired at 10 nm lateral resolution. z is the depth direction, with xy images acquired every 40 nm. **xz** and **yz** are reconstructed images depicting continuity of the image stack along the z-axis (depth). Scale bars = 10 microns.

### Image segmentation and quantification

To better visualize and quantify the cardiac ultrastructure, we segmented (delineated) from SBF-SEM images the cell nuclei, myofibrils, mitochondria, and the extracellular space. To this end, we used a combination of deep learning algorithms and tools available on the Dragonfly 4.1 software (Object Research Systems, Quebec, Canada). When independently tested against carefully annotated images (2 each from control and TOF hearts, region A), the segmentation accuracy from the deep learning algorithm was at least 90% for myofibrils, 94% for mitochondria, and 98% for nuclei. Additional manual segmentation ‘clean up’ was thus required to improve the accuracy of organelle depictions. For quantification purposes, to control the accuracy of segmentations, we used subsets of the full SBF-SEM datasets by selecting images and regions within images from the complete dataset. From selected images and image regions, we quantified the percentage of the cell occupied by nuclei, myofibrils, and mitochondria. That is, we quantified selected organelle density within cells. We found that the density of nuclei, myofibrils and mitochondria was not significantly different between the control (CON) and TOF hearts, nor between regions A and B (see **Figure 7A**). We also quantified the percentage of the image that was occupied by extracellular space. We found that the TOF heart exhibited more extracellular space than the CON heart in both regions A and B (see **Figure 7B**). This was also consistent with a visual inspection of segmented images (see **Figure 7C**). In addition, for both CON and TOF hearts, there was more extracellular space in region B than in region A (p < 0.05).

**Figure 7:**
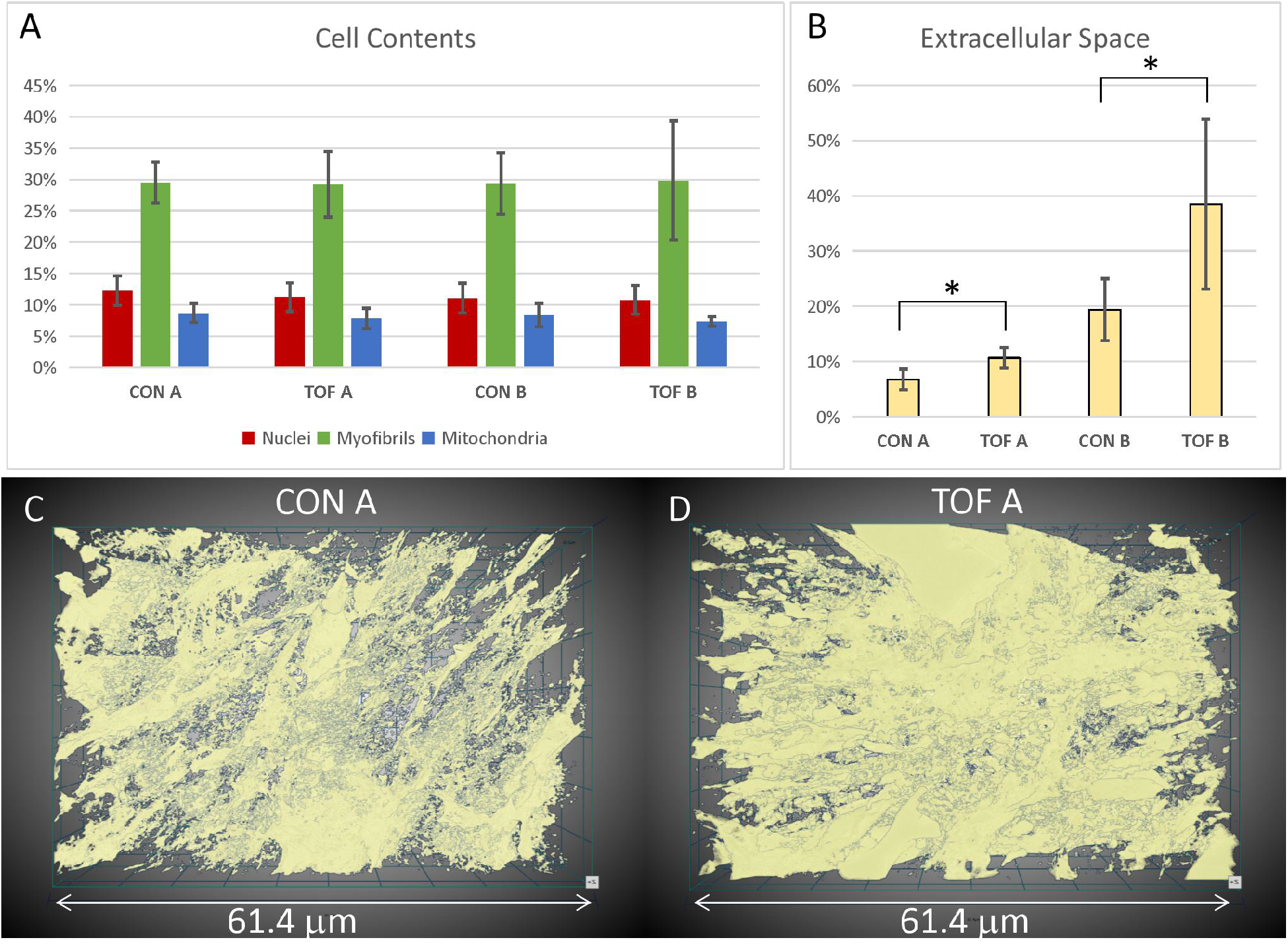
Segmentation and quantification of SBF-SEM images. Selected images (n ≥ 10) from the SBF-SEM image stacks acquired at regions A and B were carefully segmented and quantified. **(A)** Percentage of myocardial cells occupied by nuclei, myofibrils and mitochondria in two regions (A and B) of the control (CON) heart and TOF heart. **(B)** Percentage of the images occupied by extracellular space. * indicates statistically significant differences (p < 0.05). Although not marked, differences between regions A and B were also statistically significant. 3D views of the segmented extracellular space in **(C)** control (CON) heart, region A; and **(D)** TOF heart, region A.

Visualization of segmentations of the entire sub-ROIs from region A of the CON and TOF hearts revealed a slightly different orientation of myocardial cells between samples (**Figure 8**).

**Figure 8:**
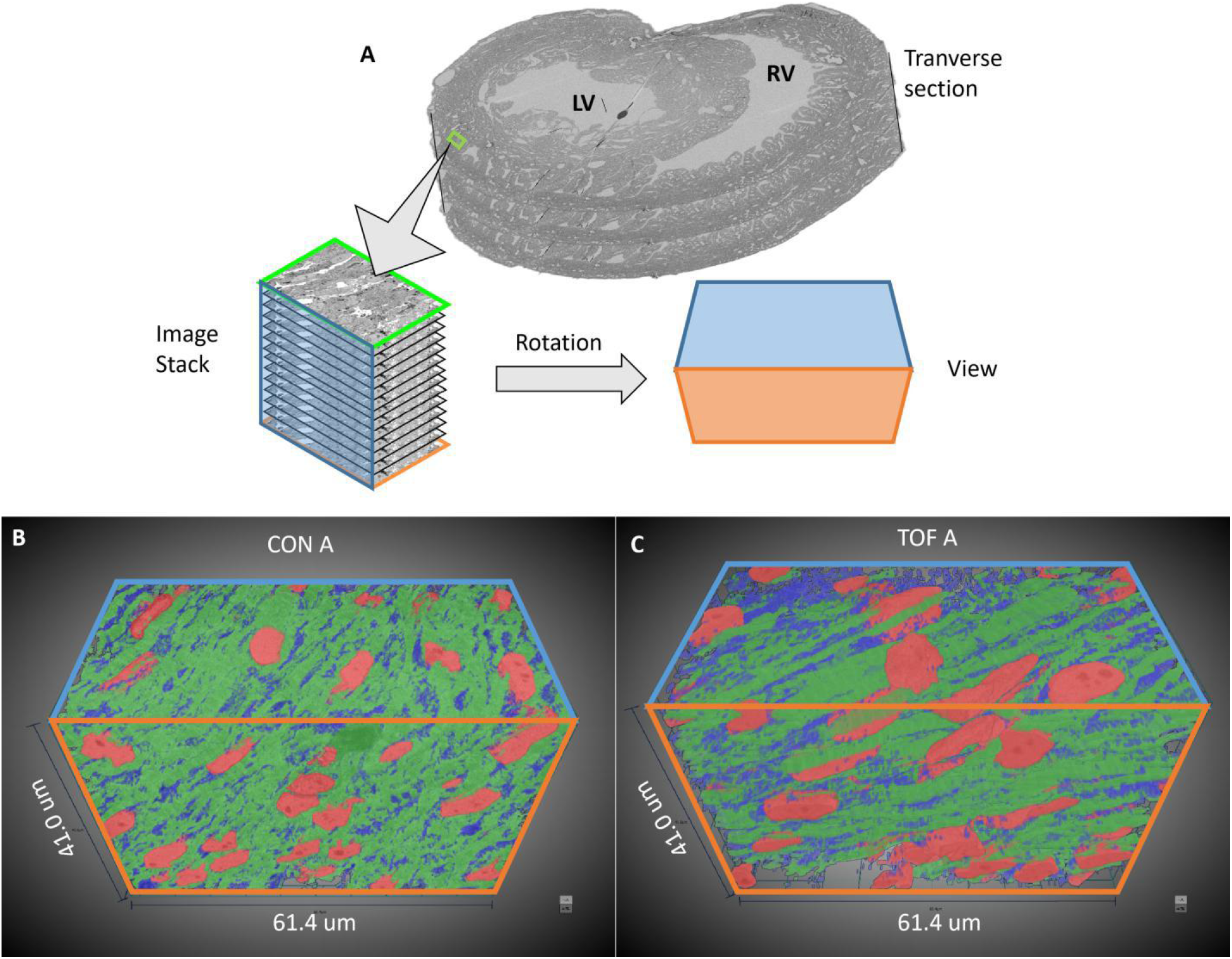
3D visualization of SBF-SEM segmentations. **(A)** Sketch of image dataset acquisition, showing relative orientations. Green plane is the top image, orange plane is the bottom image (last image acquired), blue plane is approximately parallel to the heart wall. The sketch of the view shows the orientation of the planes as shown in **(B)** control (CON) heart and **(C)** TOF heart.

## Discussion

Heart function relies on a multiscale, finely orchestrated contractile machinery. The heart ultrastructural organization is needed for efficient heart pumping and is linked to the cell metabolism^23–25^. Structural malformations in the chambers, valves or vessels of the heart, together with disruptions to the organization or number of cardiac cells, and/or their ultrastructure, can compromise the heart’s ability to pump blood efficiently^8,10,13,26,27^. Anomalous structural and ultrastructural architectures can be detrimental to the heart’s capacity to adapt to new conditions imposed by corrective surgeries or other therapies intended to repair structural congenital heart defects. Thus, the organization of the heart at multiple scales needs to be properly understood and accounted for in CHD treatment planning.

Heart walls contain myocardial cells, which are elongated, cylindrical-like muscle cells that are aligned in patterns that optimize cardiac contractility^28,29^. Myocardial cells are about 50-150μm long and 10-20μm thick (diameter), and in the heart are connected to each other forming a 3D network^11^. More specifically, myocardial cells are organized in laminar sheets that exhibit characteristic cell orientations (elliptical and transmural angles), which change over the heart wall thickness and the cardiac cycle^11,30^. At the ultrastructural level, myocardial cells contain contractile units, the myofibrils, which are supplied with energy (ATP) by the mitochondria surrounding them. In a healthy, mature heart, myocardial nuclei have an ellipsoidal shape aligned with the cell long axis; myofibrils are meticulously aligned and organized along the myocardial cells; and mitochondria are densely packed around the myofibrils^16,29^. Other organelles (such as lipid droplets and glycosomes) are organized around the cell myofibrils, mitochondria and nuclei^31,32^. While this organization may vary significantly from individual to individual, even in normal hearts, it ensures proper cardiac function. Multiscale studies of the heart, spanning whole organ to ultrastructural details, can reveal subtle deficiencies in CHD heart tissues, and relationships between abnormalities across spatial scales.

The correlative, multiscale imaging approach presented here was implemented and optimized in a chicken embryo model of heart development and CHD^33^. Heart dimensions ranged from approximately 5-6 mm long, 3-4 mm wide, and 250-700 μm wall thickness. We acquired images of the whole embryonic heart using 3D micro-CT and of cardiac ultrastructure using 3D SBF-SEM. Our multiscale imaging procedure achieved high-resolution images exhibiting both microstructural and ultrastructural preservation, while protocol completion was achieved in about 4 days, which is comparable to completion timings for much smaller samples^34^. Our multiscale imaging methodology will enable studies of yet unknown tissue deficiencies in CHD.

### Our protocol in relation to previous works

Researchers have used EM techniques, including SEM, for decades to visualize the organization of organelles within cells^20,35^. In the heart, studies using EM have revealed the ultrastructural architecture of mature myocardial cells^16,31^, and elucidated the maturation of myocardial ultrastructure during embryonic development^28,36,37^. A few studies, moreover, have determined changes in ultrastructure due to pathophysiological conditions in the mature heart^13,26,27^. Properly preparing samples for EM requires meticulous protocols that aim at preserving the ultrastructure of the tissue under study (e.g. intact cell and nucleus membranes, mitochondria and their crystae, myofibrils and z-disks, etc). Because the ultrastructural features analyzed are at the nanometer scale, samples used for EM are typically very small (< 1mm^3^), which facilitates proper sample preparation. In preparing samples, portions of the heart are typically excised, carefully prepared for imaging (fixed and stained)^14,38^, and then imaged with an EM modality^14,31^. Using this procedure, however, finding the ultrastructure associated with a specific microscopic feature can be daunting. Moreover, in micro-dissecting tissue samples, the myocardial organization within the heart may be lost^11^. Our methodology enables correlative microscopy in a way that allows precise identification and mapping of portions of the heart to their ultrastructure.

Whole animals and organs have been scanned with computed tomography (CT), a 3D x-ray imaging modality, to determine the internal and external structure of organs, including the heart. For small animal models, micro-CT, a high-resolution CT, is typically employed^33,39^. To enhance contrast and thus resolution, prior to micro-CT imaging excised tissue samples are frequently stained^33^. Preparing samples for micro-CT imaging requires good and uniform penetration of the stain, intended to preserve and contrast the tissue microstructure (e.g., heart morphology, heart chambers and valves). Micro-CT can then reveal subtle and overt malformations in the heart and its microstructure^33^. For example, using micro-CT cardiac images it is possible to visualize ventricular septal defects or translocations of the great arteries, but also wall and septum thickness, and differentiate trabecular from compact myocardium^33^. The discrepancy in the scale at which micro-CT and SBF-SEM are acquired introduces fundamental differences in the requirements for sample preparation. Due to diverse fixation and staining protocols^33,40^, tissues prepared for micro-CT or other microstructural imaging (e.g. histology) cannot typically be simultaneously processed for SBF-SEM (and other EM modalities), thereby restricting the ability to analyze both the structural and ultrastructural characteristics within the same tissue sample. Our correlative micro-CT/SBF-SEM procedure can reveal the sub-cellular architecture associated with specific pathological or malformed regions of the heart found from micro-CT images.

Applications combining micro-CT and EM technologies have recently begun to emerge, e.g.^41–43^. However, several challenges remain in applying these methods to correlative, multiscale imaging of a relatively large organ like the heart (even the heart of a small animal). To achieve both EM and micro-CT high-quality imaging of the same sample, existing protocols have capitalized on heavy metal contrast in small tissue samples, which are fully processed prior to EM and micro-CT imaging, e.g.^34,41^. However, achieving the uniform staining necessary for optimal imaging with both micro-CT and SEM becomes progressively challenging with increasing tissue sample size. This is mainly due to difficulties in achieving uniform and fast fixation (that preserves the ultrastructure), and uniform stain penetration (both for post-fixation purposes and to enhance contrast for SEM and micro-CT imaging). Modifications to the classic ROTO protocols^19–21^ for EM tissue preparation have been quite successful in achieving strong and uniform staining of relatively large samples^44,45^. However, acceptable staining was typically only up to a depth of 200 μm, and more recently 500 μm^19^ in dense brain tissues. In an attempt to stain whole brains for EM reconstruction of synapses, Mikula et al. developed the brain-wide reduced-osmium staining with pyrogallol-mediated amplification (BROPA) protocol^46^ for 3D SBF-SEM (no other heavy metals were used). While the protocol is compatible with both micro-CT and 3D SBF-SEM imaging, preparing a whole mouse brain (about 8-10 mm in diameter) using BROPA required 2-3 months. A fast BROPA protocol (fBROPA) was later developed and used to prepare whole brains from zebrafish in about 4 days^34^. Zebrafish brains, however, are significantly smaller than mouse brains (diameter of 1.1 mm vs 8-10 mm, respectively). For our hearts, we needed to achieve stain penetration of a relatively large sample (5-6mm long, 3-4 mm wide) and a fast preparation protocol was also desired. We found that preparing the heart for SBF-SEM and stopping the protocol after initial ROTO staining, was compatible with microCT and later SBF-SEM full sample processing and imaging. To our knowledge, this is the first time that correlative micro-CT/SBF-SEM imaging is applied to the heart.

### Protocol implementation

Sample preparation for SBF-SEM required strong, immediate fixation to preserve the ultrastructure of the heart tissue. We used a modified Karnovsky’s fixative with equal parts glutaraldehyde and paraformaldehyde: The paraformaldehyde rapidly penetrated and temporarily stabilized the tissue, and the slower-penetrating glutaraldehyde, a superior crosslinker, more permanently preserved the tissue sample^35^. An obvious difficulty was to achieve uniform fixation of the whole heart sample. Homogenous fixation was achieved by perfusing fixative into the heart prior to excision and then promptly immersing the heart in fixative after excision, allowing the fixative to simultaneously penetrate the heart through the tissue’s internal and external surfaces. The hearts were then post-fixed with osmium tetroxide, a lipid crosslinker, which fully stabilized membrane structures while enhancing contrast for micro-CT and SBF-SEM imaging. For our heart samples, we could achieve uniform stain penetration using a variation of the ROTO protocol with extended staining timing (about 30% increase; see Methods). The extended timing was sufficient for our hearts, even considering small size variations. We expect, however, that further time increases would apply to larger heart samples (for instance if we image embryos at a more advanced developmental stage, or other species are considered). For the chick embryos studied here, sample preparation after ROTO was adequate for micro-CT imaging of the whole heart (see **Figure 2**).

For 3D SBF-SEM images, heart samples were further stained with a combination of heavy metals. This is because SEM images are acquired via the detection of secondary and backscattered electrons that are emitted as the tissue is scanned with a high energy beam of primary electrons^47^. Soft biological tissues, like the heart muscle, yield few backscattered electrons and need to be stained with heavy metals, which readily produce secondary and backscattered electrons. Heavy metal stains interact with specific ultrastructural components, and therefore combinations of stains are frequently used in a single sample. The application of osmium tetroxide, used in ROTO protocols, served as the first application of a heavy metal stain in the SBF-SEM preparation. The osmium tetroxide, which interacts with lipids in membranes and vesicles, both post-fixed and stained tissues. Further staining with uranyl acetate stained lipids and proteins, and lead aspartate stained proteins and glycogens. Together, these heavy metal stains fully preserved and contrasted ultrastructural details, as evidenced by high-resolution SBF-SEM images (see **Fig. 5**).

For whole-heart samples, we found that staining with uranyl acetate and lead aspartate rendered resin-embedded hearts opaque to micro-CT imaging (data not shown). Our multiscale multimodality imaging procedure overcame these difficulties by performing micro-CT imaging after ROTO (see above) but prior to the lead and uranium staining steps, such that correlative 3D micro-CT and 3D SBF-SEM could be implemented. This initial modified ROTO post-fixing and staining provided tissue contrast for micro-CT while also ensuring that the sample was preserved and stabilized before final SBF-SEM sample preparation and imaging (see **Figure 1**). In addition, it enabled screening, storing and selection of hearts before final SBF-SEM processing. This feature of our multiscale approach is advantageous for several reasons. At this early processing point, hearts could be stored for relatively long periods (at least 2 weeks, but potentially months/years) before further processing. This allows researchers to prepare and screen by micro-CT a large number of hearts, and then select only those hearts of interest (e.g. with a specific malformation) for further analysis with SBF-SEM. Not only does this approach permit banking of samples, but it also saves considerable time and resources by avoiding full sample preparation of hearts that are not useful. This is important for our application, in which at most 60% of treated hearts develop structural malformations, and the nature and severity of defects vary among individual hearts. Since we cannot accurately classify malformations until they are scanned with micro-CT, being able to use the micro-CT as both a screening tool and as a navigational tool for later correlative microscopy is invaluable to CHD studies.

Due to the size of the hearts and slow diffusion (penetration) of heavy metals into the tissue, achieving uniform staining during further SBF-SEM tissue processing was not straightforward. Initial iterations of the procedure (data not shown) resulted in a strong gradient of staining into the heart tissue, a manifestation of poor stain penetration^34,46^. Application of microwave steps to enhance diffusion was not helpful, although perhaps optimizations of those steps could achieve improved results. We found, however, that following a Renovo Neural, Inc. protocol (see Methods) after ROTO and before embedding the sample in a resin capsule allowed the samples to achieve homogeneous staining. Uniform stain penetration throughout the heart, was apparent from semithin transverse sectional images of the heart and low-resolution images of ROIs (see **Figure 4**). The uniform contrast and resolution of ultrastructural details provided further evidence of uniform staining and proper tissue fixation (see **Figure 5**).

While the focus of this study was to demonstrate homogeneous staining and fixation, our procedure enables correlative microscopy. One way to achieve accurate localization of ultrastructures within the heart structure, is to register semithin transverse sections to micro-CT images, and then SBF-SEM images to the semithin images (as done in **Figure 4**). Because sectioning of samples for SBF-SEM imaging is done after micro-CT images are acquired, sectioning is guided by the images of the whole heart, facilitating the selection of regions of interest. Registration can then be performed among the images themselves. This could be done directly, or by adding fiduciary markers in the resin/heart to facilitate image alignment. Our procedure allows imaging of ultrastructure at several regions of interest within the heart, enabling extensive ultrastructural mapping.

### Comparison of control and TOF hearts and limitations of this study

We further explored some possible analysis and quantification strategies enabled by our procedure. We acknowledge that any results from this study are very preliminary, and any statistically significant differences show here pertain only to the two hearts studied. Analysis of more heart samples is needed to reach conclusions applicable to CHD. In the future, a combination of segmentation, quantification and other refined methods to interrogate the images (at the microstructural and ultrastructural levels) will elucidate similarities and differences between normal and malformed hearts, possibly informing therapeutic treatment strategies.

Micro-CT images reveal the microstructural characteristics of hearts. Not only could we classify the hearts based on phenotype (normal vs TOF), but (while not reported here) cardiac characteristics, such as heart size and wall thickness could be visualized and quantified. For example, it was noted from our analysis that the RV of the TOF heart exhibited a larger volume and thinner ventricular walls than that of the control (CON) heart examined here (see **Figure 3**). During fetal life, the lungs are not functional, and blood to the lungs is shunted to the systemic circulation through the ductus arteriosus^48^ (a pair of ducti in chick^49^). The RV hypertrophy characteristic of TOF, develops over time after the baby is born^50^. RV hypertrophy, therefore, may not be present at the fetal stages of heart development examined in this study and was not observed in our TOF heart. In the future, it would be interesting to determine whether the trait observed in this study is preserved among TOF fetal hearts, and if so under which conditions and how it affects the RV wall ultrastructure (not examined here). Such study, however, requires a larger number of heart samples, and is outside of the scope of this paper.

Another difference between the two hearts, was that the ventricles of the TOF heart exhibited less dense tissue and a more extended trabecular architecture compared with the control heart. This is consistent with reduced myocardial compaction in TOF^51–53^. The heart trabeculae is characterized as a “spongy” or porous tissue that develops inside the heart ventricles, and in our samples was evident from semithin transverse heart sections (**Figure 4**), but could also be approximately quantified as the extracellular portion of the images (**Figure 7**). It has been shown that the heart trabecular architecture is sensitive to blood flow conditions during development^54^, and thus a disrupted trabecular architecture may be a characteristic of TOF hearts due to their anomalous flow characteristics during fetal stages^18^. However, the trabecular and myocardial architecture can also exhibit variations from heart to heart^11^, therefore further analysis with a larger sample size is required before we can make conclusions related to TOF.

We noticed differences in SBF-SEM image sharpness, which are attributable to excessive ‘charging’ (accumulation of static charge on a sample’s surface) when scanning the TOF heart (**Figure 5**). The increased charging in our TOF heart is linked to its trabeculation, which features larger and more numerous void regions filled with free resin. To address this problem during imaging, we slightly shortened the dwell time when acquiring SBF-SEM images from the TOF sample. In the future, we could embed silver particles in the resin to increase sample conductivity and enhance image quality^34^. Nevertheless, the image quality of both the TOF and the control hearts was sufficient to appreciate ultrastructural details (**Figure 5**).

While 2D EM images have been invaluable in deciphering ultrastructural features of myocardial cells and tissues, 3D images can unravel more details in the spatial organization of the ultrastructural architecture^16,26^. As an example, segmentation and visualization of the 3D data revealed that myofibril alignment was slightly different between our TOF and control hearts (**Figure 8**). This is perhaps because the ROIs from the two hearts are not exactly corresponding, or due to the more extended trabecular architecture of the TOF heart, and warrants further investigation. For myocardial alignment quantification, it is also important to arrest the heart consistently (in diastole as done here, or systole) as myocardial cell orientation changes over the cardiac cycle^12,30^. 3D SBF-SEM images also revealed a greater proportion of endocardial cells in TOF heart tissues than in control tissues, such that volumetric studies not focusing on myocardial cells show reductions in the myofibril density of the TOF heart (data not shown). When the analysis is focused exclusively on myocardial cells, however, we could not find any differences in the density of myofibrils or mitochondria (**Figure 7**). While outside the scope of this paper, future studies should focus on elucidating ultrastructural cardiac differences in animal models of CHD as such differences can impact the lives of children and adults with congenital heart defects. Our proposed multiscale imaging methodology will certainly enable such studies.

## Conclusions

Our correlative, multiscale imaging procedure allowed us to acquire detailed micro-CT images of an entire embryonic chicken heart (see **Figure 2**), followed by ultrastructural 3D SBF-SEM images from the same heart (see **Figures 4** and **5**). Our approach thus enables detailed analysis of both whole heart morphology and ultrastructural architecture, allowing determination of cardiac malformations and subcellular organization within specific regions of the heart. This is important when studying CHDs, as each malformation phenotype may be different and therefore may need to be analyzed separately to fully appreciate multiscale effects and to understand how phenotypes affect cardiac architecture at disparate levels. The multiscale imaging approach presented here will enable studies to determine how cardiac anomalies, even when repaired, could subsequently lead to increased cardiac dysfunction and heart failure. For patients with CHD, such studies may reveal associated pathologies in cardiac tissues that, if not properly treated, may have devastating implications for survival and long term cardiac health.

While our developed multiscale approach was implemented and optimized using embryonic chicken hearts, we expect it will be straightforward to adapt it for use in mouse and other small animal models of cardiac malformations. It will be advantageous to use complementary models, as typically genetic insults are studied using mouse models, while environmental perturbations are studied using avian models. Heart dimensions in those species (mouse and chick) are very similar, and we anticipate that tissue processing will not differ significantly. Slight increases in heart size may just require an increase in protocol staining times. Extending the approach to different species and models of congenital heart disease will enable us to understand in detail the similarities and differences between cardiac defects, and the underpinnings of malformations that result from genetic and environmental insults.

## Methods

### Ethical Considerations

Our research used chicken embryos. According to the US National Institutes of Health (NIH) Office of Laboratory Animal Welfare (*ILAR News* 1991; 33(4):68-70), the NIH’s “Office for Protection from Research Risks has interpreted ‘live vertebrate animal’ to apply to avians only after hatching.” Our Institutional animal care and use committee (IACUC) follows NIH interpretation. Therefore, chicken embryos are not considered animals and our research did not require approval. Incubator logs in the lab were monitored daily to ensure there were no eggs near the hatching time of 21 incubation days. Nevertheless, we used the minimum possible number of embryos to achieve our goals.

### Generation of cardiac defects

Our multiscale approach was implemented and optimized using fully formed embryonic chicken hearts (heart length ~5-6mm), and applied to a chick animal model of congenital heart disease. Chicken embryos were prepared as described previously^33^. Briefly, fertilized white Leghorn chicken eggs were incubated blunt end up at 38°C and 80% humidity for approximately 3 days (to Hamburger and Hamilton (HH) stage HH18^55^). Control and treatment interventions were then performed as described below and the embryos were re-incubated for an additional 9 days (to HH38, when the heart has four chambers and valves). Two embryonic hearts were included in this study: 1) a control, normal heart; and 2) a malformed heart with tetralogy of Fallot (TOF). TOF was achieved by performing outflow tract banding (OTB) at HH18, wherein a 10-0 nylon suture was passed under the mid-section of the heart outflow tract and tied in a knot (band tightness 38%). The band was removed from the outflow tract ~24 hours after placement (HH24), and then the embryo was allowed to develop to HH38. The control heart was obtained by passing a 10-0 nylon suture under the heart outflow tract without knotting it, and subsequently allowing the embryo to develop to HH38. Embryo hearts were collected at HH38 for multiscale imaging.

### Homogenous fixation of whole hearts

At HH38, embryonic whole hearts were excised and fixed as follows. The chest cavity was opened and the pericardial sac around the heart gently removed with forceps. Each heart was arrested by injecting 500 μL of chick ringer solution containing 60 mM KCl, 0.5 mM verapamil, and 0.5 mM EGTA^56^ into the left ventricle through the heart’s apex. Hearts were then immediately perfused with ~2 mL of ice-cold (0°C) modified Karnovsky’s fixative (2.5% Glutaraldehyde and 2.5% PFA in PBS (pH 7.4)) through the same injection site. All perfusions were performed with a 21 gauge needle. A transfer pipette was used to quickly apply ~1 mL of fixative to the heart’s exterior to ensure uniform fixation of the heart tissue. Next, the heart great vessels were cut with small spring scissors and hearts were placed in 1.5 mL fixative and stored at 4°C until further processing.

### Cardiac processing enabling micro-CT imaging

In order to enable both whole-heart micro-CT imaging and subsequent SBF-SEM imaging of regions of interest (ROIs), we processed fixed hearts for micro-CT using the initial portion of a Renovo Neural, Inc (Cleveland, USA) protocol^38^ designed for SBF-SEM imaging (see **Figure 1, Step 1**). Each heart was placed in a 5 mL glass scintillation vial and we used 3 mL of solution per vial for each incubation/wash. First, the fixed hearts were washed in 0.1M Sodium Cacodylate (pH 7.4) for 20 minutes with 4 exchanges of fresh buffer. Next, the hearts were incubated in 0.1% (w/v) of tannic acid in 0.1M Sodium Cacodylate (pH 7.4) for 15 minutes at room temperature. Samples were then washed in 0.1M Sodium Cacodylate (pH 7.4) for 20 minutes with 4 exchanges of fresh buffer. Since the reducing agents used in subsequent steps (modified ROTO protocol) were light-sensitive, the sample vials were covered in aluminum foil from this point on. The whole hearts were post-fixed in 2% (v/v) Osmium Tetroxide (OsO_4_) and 1.5% (w/v) Potassium Ferricyanide (K_3_[Fe(CN)_6_]) in distilled water (dH_2_O) for 2 hours at room temperature on a rotating platform. The samples were then extensively washed in dH_2_O for 20 minutes with 4 exchanges of fresh dH_2_O. Next, the samples were immersed in 0.1% (w/v) Thiocarbohydrazide (TCH) solution in dH_2_O, placed in an oven, and incubated for 40 minutes at 60°C. This step was followed by another 4 exchanges of fresh dH_2_O over 20 minutes. Samples were then immersed in a 2 % (v/v) OsO_4_ solution in dH_2_O for 2 hours at room temperature on a rotating platform. Finally, the hearts were washed extensively in dH_2_O over 20 minutes with 4 exchanges of fresh water. Each heart was stored in dH_2_O at 4°C until imaged by micro-CT. This preparation provided excellent contrast for micro-CT scans (see Results).

Micro-CT images of whole hearts were acquired to assess the cardiac structure. We acquired high-resolution (~10 μm) 3D scans of each heart using a Caliper Quantum FX Micro-CT system (Perkin-Elmer, CLS140083) with 10 mm field of view, 140 μA current, 90 kV voltage, and a scan time of 3 minutes. We used the Amira 6.0 software platform (FEI Company) to visualize these scans and identify cardiac defects. Hearts were then stored in double distilled water at 4°C until further processing. Please note that at this step in the processing (cardiac tissues fixed and post-fixed with OsO_4_) water does not damage the tissues.

### Subsequent cardiac tissue processing enabling 3D SBF-SEM imaging

After whole hearts were imaged with micro-CT, sample preparation of selected hearts for 3D SBF-SEM imaging was finished (see **Figure 1; Step 2**), following the Renovo Neural, Inc. protocol. In large samples, like the whole hearts described in this manuscript, it is necessary to extend most of the staining steps. Failure to extend the timing of staining resulted in a heterogenous stain distribution throughout the tissue (in our early iterations of the procedure). In our final, optimized procedure, we incubated the samples in 1% (w/v) aqueous uranyl acetate for 24 hours at 4°C, after which they were washed in dH_2_O for 30 minutes with 6 exchanges of fresh dH_2_O. We then incubated the samples in lead aspartate for 30 minutes at 60°C. The samples were then extensively washed in dH_2_O for 20 minutes with 4 exchanges of dH_2_O. Dehydration steps were done in a series of acetone-dH_2_O mixtures (50, 75, 85, 95 and 100%); each step was repeated twice for 5 minutes at room temperature. The whole heart sample was then embedded in an epoxy (Epon 812) resin for further manipulation and SBF-SEM sample preparation. The first infiltration step was done for 1 hour at room temperature in a mixture of 1:1 (v/v) acetone:epon followed by a 1:3 (v/v) acetone:epon incubation for 1 hour at room temperature. The hearts were subsequently incubated overnight in pure (100%) epon on a rotating platform. The following day the epoxy solution was exchanged 4 times, each time with 30 minutes incubation steps at room temperature. Samples were polymerized at 60°C for 48 hours in a conventional oven, leading to a whole heart sample embedded in an Epon block.

### Selection of regions of interest (ROIs) and SBF-SEM image acquisition

Using the micro-CT images as reference, the Epon-embedded heart blocks were sectioned to reach a selected short axis (transverse) section using a diamond-wire jewelry saw. For this study we selected the mid cardiac transverse section, at a plane where the heart is wider (the equatorial plane). After this step, a semithin section (250 nm) was obtained using an ultramicrotome and mounted on a silicon chip previously glow discharged for 1 minute at 15 mA (PELCO easyGlow, Ted Pella). Semithin section images were used to confirm the area of interest as well as to check for both the ultrastructural quality of the sample and the success of the staining procedure (see **Figure 1; Step 2**). This step is crucial since the SBF-SEM imaging requires samples with extremely good contrast. Semithin sections were imaged on a Teneo Volume Scope in low vacuum mode using a VS-DBS backscattered electron detector and the MAPS software (FEI Company). Imaging conditions used were 2.5 kV and 0.2 mA, dwell 3-5 μs. In some cases, the samples imaged using this method needed to be coated with a thin (5-8 nm) layer of carbon to minimize charging artifacts induced by the electron beam.

The same diamond-wire jewelry saw was then utilized to generate a slab (~1.5 mm) from the sample (see **Figure 1; Step 3**). The slabs were sectioned into smaller ROIs, which were subsequently mounted on Microtome stub SEM pins (Agar Scientific 61092450) using H20E Epo-Tek silver epoxy (Ted Pella 16014) and cured overnight at 60°C in a conventional oven. The resulting small blocks were then trimmed using a Trim90 diamond knife (Diatome) to generate a pillar of 500×500 μm^2^. The block was then coated with 20 nm of gold using a Leica ACE 600 unit.

In the last step of our multiscale imaging procedure, 3D SBF-SEM images of sub-ROIs selected from the mounted sample were acquired (see **Figure 1; Step 4**). 3D image acquisition was done on a Teneo Volume Scope SBF-SEM in low vacuum mode (50 Pa) using a VS-DBS backscattered detector. Images were acquired at a lateral resolution of 10 nm/pixel and image sets included 800-1000 serial sections (with each section thickness measuring 40 nm in the z axis). SBF-SEM data sets were approximately 40 μm × 60 μm × 32-40 μm.

### Image Analysis and Segmentation

All registration, of SBF-SEM data was performed with Amira 6.0 (FEI Company). First, complete image stacks (800-1000 slices) from each ROI and sub-ROIs were automatically aligned to generate a continuous 3D volume. Next, a non-local means filter was applied to every 2D slice in order to improve the signal-to-noise ratio. Due to slight differences in the intrinsic properties of the tissue, sections from the TOF heart appeared slightly lighter compared to the control heart. We adjusted the intensity of the TOF sections during post-processing to match that of the control heart.

To better appreciate ultrastructural differences between the two hearts, we used Dragonfly 4.1 software (Object Research Systems, Quebec, Canada) to segment and quantify SBF-SEM images. We segmented: cell nuclei, mitochondria, myofibrils and the extracellular space (this later one to allow quantification of relative organelle volume within cells). The segmentation used a combination of tools in Dragonfly, including deep learning algorithms. Briefly, we employed a six-level U-Net deep learning model^57^ implemented in Dragonfly to perform an initial segmentation of nuclei, mitochondria, and myofibrils within imaged cells. Training sets required for the deep learning model were obtained initially through manual segmentations of a few selected images from the image stacks. The training set was later augmented by applying the segmentation deep learning algorithms to other (selected) images from the set, follow by thorough manual cleaning. For each training session, the model was run for 50 epochs with a patch size of 128 pixels. Dragonfly automatically divides the training sets into training and validation regions, so that training (and further inclusion of training images) continued until the reported accuracy (from validation regions) was > 98% with a loss < 0.06. Images were then segmented with the trained model, and segmentations further refined both using Dragonfly automated tools, such as morphological operations, and manual clean up using painter tools available in Dragonfly. Unlike organelles, the extracellular space was easily recognized by intensity levels, and thus simply segmented based on its intensity, followed by clean up using both automatic and manual tools in Dragonfly.

For visualization purposes, we segmented a whole dataset from the control heart and an approximately corresponding dataset from the TOF heart (region A, **Figure 8**). The total volume of the dataset was 40×60×32 μm^3^. For quantification purposes, we segmented and further curated smaller portions of the data sets from two corresponding regions of the control and the TOF hearts (regions A and B). Quantifications of extracellular space and nuclei were done from 17 evenly spaced images from the 800 image datasets of region A; and 21 evenly spaced images from the 1000 image datasets of region B (thus every 50^th^ image was used for these segmentations). For quantification of mitochondria and myofibrils, we cropped images (n ≥ 10) so that we could focus on smaller regions, allowing us to manually improve the accuracy of segmentations in a more tractable manner and focusing on myocardial regions.

Quantifications were performed based on segmented images. We quantified, from each image or image portion, the total surface area (S_T_), and the surface area occupied by extracellular space (S_E_), nuclei (S_N_), mitochondria (SMit) and myofibrils (SMyo). We then computed the fraction of the total surface area occupied by the extracellular space (S_E_/S_T_); and the fraction of the cell occupied by organelles (nuclei, mitochondria and myofibrils), computed as the ratio of organelle surface (S_i_, with i = N, Mit, Myo) to the cell surface (S_i_/(S_T_-S_E_)). Because quantifications were performed from different portions (n ≥ 10) of the dataset, average and standard deviations were calculated to represent quantifications for the dataset (control or TOF hearts, regions A and B). We then employed a one tail T-test to compare quantifications among datasets, with p < 0.05 indicating significant differences.

## Acknowledgements

This work has been funded by a grant from US National Institutes of Health, NIH R01 HL094570 (SR); the OHSU University Shared Resource pilot funding program (SR); the OHSU School of Medicine Faculty Innovation Fund (SR). We would like to thank Kevin Loftis for graciously sharing his time and knowledge of Amira. We would also like to thank Melissa Williams for her help in optimizing our sample preparation and Renovo Neural, Inc, especially Emily Benson and Grahame Kidd, for sharing their SBF-SEM protocol. Electron microscopy was performed at the OHSU Multiscale Microscopy Core (MMC) with technical support from the OHSU Center for Spatial Systems Biomedicine (OCSSB).

An extended abstract (2 pages) of this work was published and presented at Microscopy & Microanalysis 2019 Meeting in Portland, OR, https://www.cambridge.org/core/journals/microscopy-and-microanalysis/article/multiscale-cardiac-imaging-from-whole-heart-images-to-cardiac-ultrastructure/7053ED929882C2E43B2ED85FD6D78BEC.

## Author Contributions

G.R. prepared the samples, acquired micro-CT images, and analyzed micro-CT and EM images. C.S.L and J.L.R. optimized the sample preparation protocols and acquired the EM images. I.F. segmented and quantified the SBF-SEM images. S.D. segmented micro-CT images for comparison. K.C. analyzed images and assisted G.R. with figures. A.M and K.T. analyzed the quality of images and helped with biological image interpretation. S.R. conceived the project, and wrote the first draft of the manuscript. All authors reviewed and edited the manuscript.

## Data Availability

Data, including micro-CT and SBF-SEM images, will be available to researchers upon request. Protocols employed are fully disclosed and detailed in the manuscript methods section.

